# An *ex vivo* cystic fibrosis model recapitulates key clinical aspects of chronic *Staphylococcus aureus* infection

**DOI:** 10.1101/604363

**Authors:** Esther Sweeney, Marwa M. Hassan, Niamh E. Harrington, Alan R. Smyth, Matthew N. Hurley, María Ángeles Tormo-Mas, Freya Harrison

## Abstract

*Staphylococcus aureus* is one of the most prevalent organisms isolated from the airways of people with cystic fibrosis (CF), predominantly early in life. Yet its role in the pathology of lung disease is poorly understood. Clinical studies are limited in scope by age and health of participants and *in vitro* studies are not always able to accurately recapitulate chronic disease characteristics such as the development of small colony variants. Further, animal models also do not fully represent features of clinical disease: in particular, mice are not readily colonized by *S. aureus* and when infection is established it leads to the formation of abscesses, a phenomenon almost never observed in the human CF lung. Here, we present details of the development of an existing *ex vivo* pig lung model of CF infection to investigate the growth of *S. aureus*. We show that *S. aureus* is able to establish infection and demonstrates clinically significant characteristics including small colony variant phenotype, increased antibiotic tolerance and preferential localisation in mucus. Tissue invasion and the formation of abscesses were not observed, in line with clinical data.

## Introduction

*Staphylococcus aureus* is one of the most prevalent pathogens isolated from the airways of people with cystic fibrosis (CF) (1). It is predominantly associated with paediatric pulmonary infection, particularly in the first decade of life (2). The presence of *S. aureus* in the respiratory tract varies considerably both geographically and over time, but prevalence does appear to reduce with age (3). Determining the difference between colonisation and infection is both important and difficult. Nasal carriage of *S. aureus* among children with cystic fibrosis is common: Stone *et al.* (4) reported that 52.4% of the infants they studied harboured the organism. However, relatively high carriage rates in healthy children have also been recorded (5). Further, expectorate is difficult to collect from infants and so samples are usually collected by oropharyngeal swab. This creates problems interpreting the presence of organisms in samples, because presence in the upper respiratory tract is not always indicative of lower airway infection (6, 7).

Moreover, the association of *S. aureus* with progressive lung disease – as measured by worsening lung function and the development of subsequent infection by the chronic CF pathogen *Pseudomonas aeruginosa* – is unclear (8). *S. aureus* is able to rapidly adapt to and persist in the CF lung environment (9), and worsening lung condition has been associated with the formation of *S. aureus* small colony variants (SCVs) (10). SCVs are known to remain in the lung longer than wild type (WT) bacterial cells (11) and demonstrate increased antimicrobial resistance (12). In studies of bronchoalveolar lavage (BAL) fluid in children aged 0-7 years, both Gangell *et al.* and Sagel *et al.* (13, 14) found that positive *S. aureus* culture was linked to a higher degree of airway inflammation as measured by increased neutrophil count and IL-8. In another study, lung damage (bronchiectasis) in early CF was recorded by CT scan and *S. aureus* was the most commonly isolated organism (15). Conversely, in adult CF patients, *S. aureus* infections, in the absence of *P. aeruginosa,* are a marker of milder disease (1). There is also an indication that methicillin sensitive *S. aureus* may inhibit *P. aeruginosa*, thus delaying lung disease progression (16). The evidence that early *S. aureus* infection worsens prognosis for CF patients remains conflicting and warrants further study (3)(17).

The lack of understanding regarding the role of *S. aureus* in the development of lung disease in CF has led to debate over the use of anti-staphylococcal prophylaxis early in life. Studies by Ratjen *et al.* and Stutman *et al.* (18, 19) linked the use of broad-spectrum antibiotics to an increase in *P. aeruginosa* isolation. However, a recent Cochrane review (20) showed no effect of anti-staphylococcal prophylaxis on *P. aeruginosa* colonisation at 3-4 years. There was a suggested trend towards higher rates of *P. aeruginosa* at 4-6 years but as the studies reviewed did not last more than six years, conclusions about the long-term effects of prophylaxis could not be drawn. A pragmatic randomized controlled trial is currently in progress (http://www.cfstart.org.uk).

In addition to the lack of clarity over the clinical consequences of *S. aureus* colonisation, there is also a gap in our mechanistic understanding of the microbiology of *S. aureus* in CF. The interaction of *S. aureus* with the pulmonary airway in CF, and its subsequent role in pathogenesis, is not clearly documented. CF epithelial cells have been shown to have an increased abundance of aGM1, a receptor that binds *S. aureus* and *P. aeruginosa*, compared with wild-type epithelia (21). In addition, an *in vitro* study by Schwab *et al.* demonstrated that bacterial adherence to a bronchial epithelial cell line was significantly greater for CF *S. aureus* isolates than non-CF (22). Yet, the only study we could find which included direct evidence of *S. aureus* infection in human biopsy specimens, demonstrated that *S. aureus* did not adhere to the airway epithelium but was found aggregating within the mucus (23). Furthermore, it suggested that binding to mucin in CF airways may be more efficient than to mucins produced in other infections. This is supported by Yildirim *et al.* (24), who showed *S. aureus* isolated in only 10% of samples in a study of chronic sinusitis. It is obvious that better understanding of just how *S. aureus* colonizes the pulmonary airway in CF is required, as well as more unambiguous data on the underlying mechanisms of pathogenesis and virulence, and the influence *S. aureus* has on subsequent infection by other microorganisms. The development of a clinically relevant, high throughput model of infection to dissect the microbiology of *S. aureus* in CF is therefore needed.

Mice are the most commonly used animal model of pulmonary infection in CF. Mouse models have been used to identify virulence-related genes in pathogens and to test novel therapeutic agents aimed at reducing inflammation or infection (25). However, mouse models present a number of challenges. In particular, mice do not develop spontaneous *P. aeruginosa* endobronchial infection, suppurative lung disease and mucus plugging of the airways that are fundamental characteristics of human CF progression (26). *S. aureus* does not appear to readily colonize the airways of mice or produce an inflammatory response, even when clinical strains are used, in the presence of mucus (27). Furthermore, despite one murine study demonstrating that *S. aureus* did increase the likelihood of subsequent chronic *P. aeruginosa* infection, it also promoted severe abscess-like lesions in the lung (28). When lung abscess does occur in humans, *S. aureus* is the most commonly isolated organism (29). However, lung abscess is rare in children (30) and even more so in CF patients: since the advent of neonatal screening and survival past infancy, abscesses are almost never observed in the CF population (31-33).

We have previously described an *ex vivo* pig lung model of *P. aeruginosa* airway infection in CF using alveolar and bronchiolar tissue sections and an artificial sputum media (ASM) designed to mimic CF mucus (34, 35). Here, we report that the model supports the growth of *S. aureus* as biofilm-like aggregates on the surface of bronchioles and in the surrounding artificial sputum, without the formation of abscesses or other significant ultrastructural changes to tissue. We show that *S. aureus* can be maintained in the model over 7 days (a notable improvement over the short-term nature of live murine models) and appears to grow in a similar way to that observed in human disease, including the presence of SCVs and a potential preference for aggregation in the surrounding mucus rather than tissue invasion. The work presented here is of importance not only for studying the colonisation of CF airway with *S. aureus* and the progression of *S. aureus* infection, but also lays the groundwork for future studies using this model to investigate interaction between *S. aureus* and other important CF pathogens.

## Materials and Methods

### Bacterial strains

*S. aureus* Newman and USA300 Los Angeles clone (LAC) were used as examples of well documented laboratory strains. 10 *S. aureus* stains isolated form 7 CF patients at Hospital Universitari i Politecnic La Fe were included as exemplars of clinical strains. Ethical clearance was sought and obtained from the Biomedical Research Committee of Research Institute la Fe (reference 2104/0563). We chose isolates that represented a range of patient clinical presentation and bacterial phenotypes (e.g. weak or strong biofilm formation as measured in *in vitro* attachment assays, isolates from patients during periods of stable infection and episodes of acute exacerbation).

### Media and culture conditions for growth in the *ex vivo* pig lung (EVPL) model

For use in the lung model, bacterial stocks were grown overnight at 37 °C on lysogeny broth (LB) agar. Artificial sputum medium (ASM) was prepared according to Palmer *et al.* (36) with the modification that we removed glucose and supplemented with 20 μg ml^-1^ ampicillin. Our previous work suggested that glucose facilitated the growth of endogenous bacteria present in the lungs and that ampicillin helped to limit the growth of any resident bacteria in the lung that remained after sterilisation (35). We selected a concentration of ampicillin that provided the best possible coverage against endogenous populations but was sub-inhibitory for the *S. aureus* strains used (data not shown).

The EVPL model was adapted from our group’s previous work (34, 35), which in turn built on use of pig lungs for non-CF studies (37). Briefly, lungs were collected from a local butcher (Steve Quigley and Sons, Cubbington, Warwickshire) as soon as possible following abattoir delivery and processed immediately upon arrival at the laboratory. Previous antibiotic administration history of the pigs is not known, but use of antibiotics as growth promoters is banned in the EU, so use is restricted to prophylactic or metaphylactic when infection is suspected. Approximately 5mm^2^ sections of bronchiolar tissue were dissected from the lung under sterile conditions. During dissection the sections were washed three times with a 1:1 mix of RPMI 1640 and Dulbecco’s modified Eagle medium (DMEM) (Sigma-Aldrich), supplemented with 50 μg ml^-1^ ampicillin, and then rinsed once in sterile ASM without supplementation. Bronchiolar sections were transferred to a clean petri dish and sterilized under a UV lamp for 5 minutes. Previous work has shown that tissue damage as a result of UV sterilisation is minimal (not visible using light microscopy) (34). Sections were transferred to 24-well tissue culture plates: each well contained 400 μl ASM supplemented with 0.8 % w/v agarose to form a soft pad. Tissues were inoculated with the appropriate strain of bacteria using a sterile hypodermic needle (29G). The tip of the needle was lightly touched to the surface of the chosen *S. aureus* colony, and then used to prick the bronchiolar tissue for inoculation. Uninfected controls were pricked with a sterile needle for mock infection. 500 μl of ASM 20 μg ml^-1^ ampicillin was added to each well and the plate was sealed with a Breath-Easier membrane (Diversified Biotech). Plates were incubated at 37 °C for up to 7 days and refreshed with 300 μl of ASM 20 μg ml^-1^ ampicillin at 48 h if appropriate. Following incubation, tissue was rinsed in 1 ml phosphate-buffered saline to remove loosely adhering cells and processed for colony count and microscopy.

### Bacterial load assay

Tissue sections used to assay total bacterial numbers in tissue-associated biofilm were homogenized individually in 1ml phosphate-buffered saline in reinforced metal bead tubes (Fisherbrand) using a FASTPREP 24 5G homogenizer (MP Biomedicals) for 40 sec at 4.0m sec^-1^. Homogenates were serially diluted and aliquots were plated on LB and mannitol salt (MSA) agar (Oxoid, Thermo Scientific) to obtain single colonies. LB agar was used to check for contamination and measure total bacterial load, including endogenous populations. MSA agar is selective for *Staphylococcus* and was used to determine colony numbers of test strains. When bacterial load in ASM surrounding the lung tissue was measured, 300 μl of ASM was taken from each well at the same time as tissue processing and transferred to a sterile Eppendorf tube. Samples were vortexed, serially diluted and plated in the same way as tissue homogenate.

LB plates were incubated aerobically at 37 °C for 18-24 h and MSA plates were incubated in at 37 °C 5% CO_2_ for 24 h and then for an additional 48 h at room temperature in ambient CO_2_ to check for the presence of small colony variants. Total bacterial load was calculated from tissue and ASM colony numbers.

#### Microscopy

Tissue sections intended for hematoxylin and eosin staining were placed in individual tissue processing cassettes (Simport) and fixed overnight in 10% neutral buffered formalin (NBF) at approximately 20 times w/v. Sections were prepared for histopathology by an external service (University of Manchester). If samples required storage before transport, they were placed in 70% ethanol and kept at 4°C for up to 7 days. Sections were transported in PBS with residual NBF. In addition, 10 μl samples were taken from the ASM surrounding lung tissue, diluted 1:10 in PBS and 5 μl drops prepared for gram stain.

Microscopy was conducted with an Axio scope.A1 light microscope (Carl Zeiss) with AxioCam ERc5S digital camera and images processed in Zen Pro 2.1 Blue edition.

#### Colony identification

Colonies that were able to grow on MSA plates and ferment mannitol (indicated by yellow pigmentation of colony and surrounding clear zone) were regarded as *S. aureus*. Colony morphology was recorded by digital photograph. The identity of colonies that were weak fermenters or small in size was initially confirmed using hydrogen peroxide to test for the production of catalase and a staphylase assay (Oxoid). If SCVs were suspected, colonies were recovered by aerobic incubation overnight on Columbia Blood Agar and morphology was compared to the originally inoculated parent strain. PCR of large and small colonies from MSA plates was used to verify taxon identity in two ways: first, we ran colony PCR using *Staphylococcus-*specific primers (38) and once genus was confirmed we amplified the 16S-23S rRNA intergenic spacer region and sequencing was used to confirm that SCVs were *S. aureus* (primers 5’-TGCCAAGGCATCCACCG-3’ and 5’-GGCTGGATCACCTCCTT-3’).

#### Determination of rifampicin minimum inhibitory concentration (MIC)

The resistance profile of selected bacterial isolates was determined by the broth microdilution method according to the EUCAST Jan 2019 guidelines (39). Briefly, fresh bacterial inocula of selected strains were prepared at 10^6^ cfu mL^-1^. 50 μL of each strain inoculum was added to 50 μL of a two-fold serial dilution of rifampicin (Fisher Scientific) at a final concentration range 64-0.000975 μg ml^-1^ in Mueller Hinton broth (MHB), in 96-well plates (Corning Costar). Plates were incubated for 18 h at 37°C, then MIC values were determined.

#### Antibiotic tolerance in the EVPL model

Infected lung bronchioles were processed as described above for bacterial load determination and incubated for 48 h. Replica tissues were either homogenized and plated for counting of bacterial load, or washed in PBS then placed in 300 μL of rifampicin at 2 μg mL^-1^ in MHB in 48-well plates, and incubated for 24 h at 37 °C. Rifampicin-treated tissues were then washed in PBS to prevent any antibiotic carryover, homogenized and plated as described above. The bacterial load in non-antibiotic and antibiotic treated tissues was then compared. Exposing the tissue to rifampicin in this way roughly mimics nebulized administration, rather than oral/intravenous dosing in which the antibiotics would effectively be administered across the bronchial wall.

#### Statistical analysis

All data were analyzed using RStudio (Mac OS X 10.6+ version 1.0.153) (RStudio, https://www.rstudio.com) using linear models followed by ANOVA and post-hoc Tukey HSD tests where appropriate, or using Kruskall-Wallis tests, as specified in the text. The repeatability of data was analyzed using Intraclass Correlation Coefficient (ICC).

## Results

### Clinical strains of *Staphylococcus aureus* are able to establish in EPVL and are maintained for seven days

Seven clinical isolates from patients with CF were inoculated into lung samples, and growth was compared to that of the Los Angeles clone of USA300, an MRSA isolate known to form strong biofilm (Figure 1). Clinical isolates were chosen from adults and paediatric patients sampled during periods of stable chronic infection, i.e. when they did not demonstrate acute pulmonary exacerbation. These included both MRSA and MSSA, biofilm and non-biofilm formers and samples isolated alone, in the presence of *Pseudomonas aeruginosa* or with multiple additional microorganisms (Table 1). The median total bacterial load at 48 hours was 8.2 x 10^7^ CFU and at least 1 x 10^7^ CFU was recovered for all isolates with the exception of FQ142. Within one lung, results at 48 h were reproducible, (ICC = 0.87, 95% CI 0.65-0.97) but reproducibility has drastically declined by day 7 (ICC = 0.42, 95% CI -0.01-0.8). For all strains, bacterial load recovered was significantly lower at day 7 compared to 48 h (ANOVA, F_1,32_ = 73.8, *p* <0.001; mean bacterial numbers recovered had dropped to 7 x 10^4^ CFU). The bacterial load for isolate FQ142 was below the limit of detection at day 7. However, it is notable that cells were still recovered after prolonged incubation to 7 days, and the drop in numbers between days 2 and 7 was strain dependent (ANOVA, interaction F_7,32_ = 3.65, *p* = 0.005).

**Table 1.** Clinical *Staphylococcus aureus* strains isolated from sputum samples from individuals with CF. Samples and health data collected and donated by the Instituto de Investigación Sanitaria La Fe, Valencia. Sample numbers are designated as supplied.

| Sample | Patient age | Exacerbation | Biofilm | MRSA | Other microorganisms isolated |
| --- | --- | --- | --- | --- | --- |
| FQ16 | 21 | No | Strong | No | *Pseudomonas aeruginosa*,<br>*Aspergillus fumigatus* |
| FQ22 | 17 | No | Moderate | No | no |
| FQ91 | 7 | No | Weak | No | no |
| FQ128 | 20 | No | Strong | No | *Candida albicans* |
| FQ140 | 8 | No | Moderate | Yes | *Pseudomonas aeruginosa*,<br>*Candida albicans* |
| FQ142 | 13 | No | None | No | *Candida albicans*,<br>*Aspergillus fumigatus* |
| FQ151 | 8 | No | Moderate | Yes | no |
| FQ184 | 9 | Severe | Moderate | Yes | *Pseudomonas aeruginosa* |
| FQ203 | 22 | No | Moderate | No | *Pseudomonas aeruginosa*,<br>*Burkholderia cepacia* |
| FQ210 | 9 | No | None | Yes | *Pseudomonas aeruginosa*,<br>*Candida albicans*,<br>*Haemophilus influenzae* |

**Figure 1.**
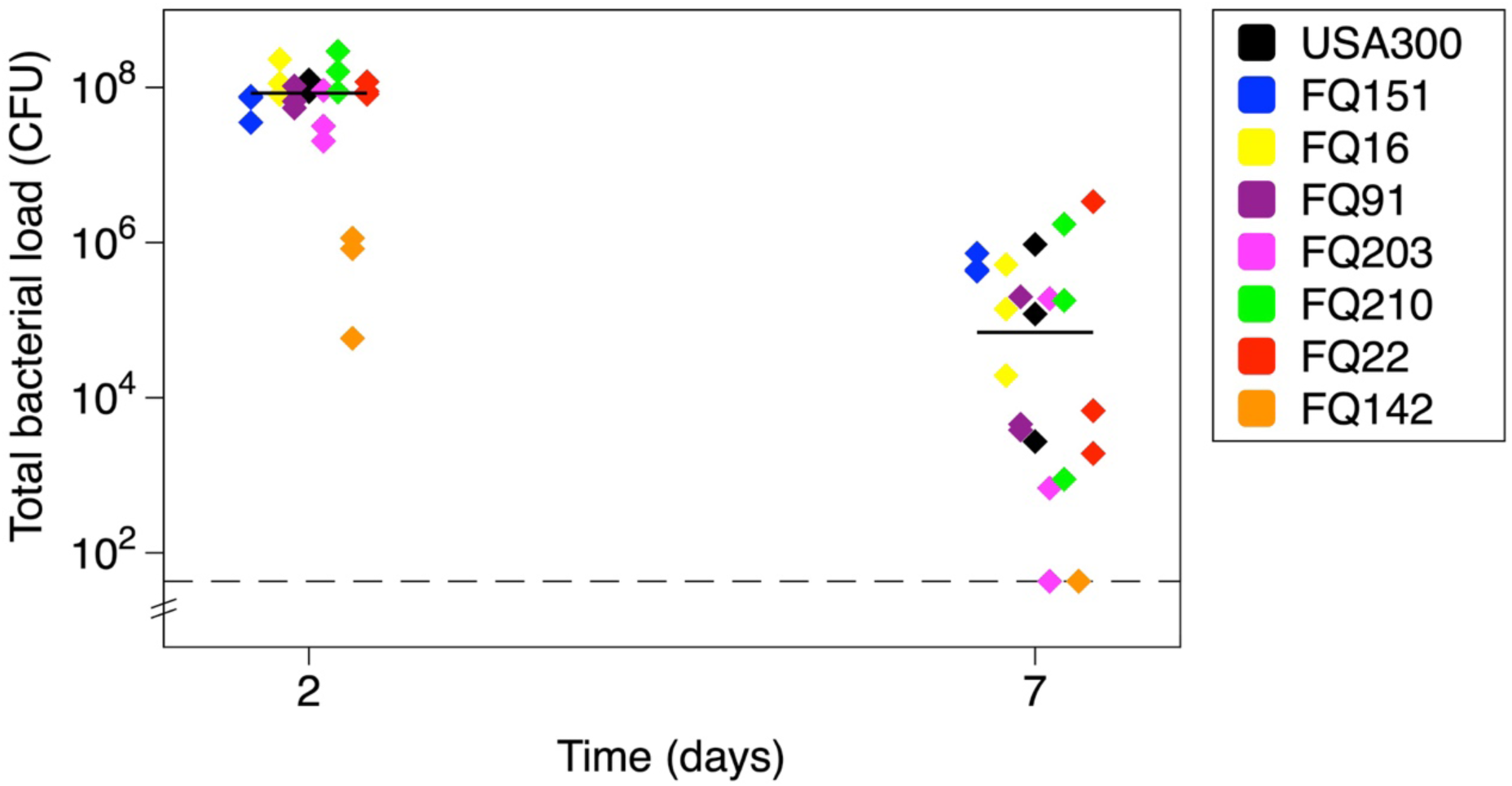
Growth of clinical *Staphylococcus aureus* over 7 days in *ex vivo* pig lung model (EVPL). *S. aureus* isolates were inoculated into six replica bronchiolar sections of tissue (all from a single pair of lungs) and samples were destructively sampled at 48 h and 7 days post infection. Bacterial load is measured as total colony forming units per lung sample (Total CFU). Lines indicate median values and the limit of detection is shown as a dashed line. USA300 is used as a representative control strain known to be a strong biofilm former. Clinical isolates were supplied by the Instituto de Investigación Sanitaria La Fe and numbers are designated as supplied. Analysis of untransformed by ANOVA showed statistically significant differences in the bacterial load attained by the different strains (F_7,32_ = 3.70, *p* = 0.005) and an overall difference in bacterial loads between 48 h and 7 days (F_1,32_ = 73.8, *p* < 0.001), as well a strain-specific pattern of the drop in CFU by day 7 (interaction F_7,32_ = 3.65, *p* = 0.005).

### Growth within the model depends on strain and lung but trends in growth are consistent overtime

To further test the reproducibility of the results for the model, and the ability to observe differences between strains within the model, triplicate samples of USA300 and two clinical strains were inoculated into three further lungs taken from different pigs (Fig 2). Clinical isolate FQ151 was compared to FQ184, a strain demonstrating similar characteristics to FQ151 but isolated during an acute disease exacerbation (Table 1). Figure 2 shows that, when repeated, USA300 was again able to establish in the lung and mean bacterial load recovered at 48 h was 6.8 x 10^6^ CFU. Mean yields for clinical strains at 48 h were 3.8 x 10^6^ and 5.5 x10^5^ CFU for FQ184 and FQ151, respectively. By 7 days no colonies were recovered from tissue taken from one of the three lungs for either clinical strain, and counts for clinical strains in a second lung were below or close to the lower limit of detection. Average CFU counts at day 7 were 4.2 x 10^3^ for USA300, 7.9 x 10^3^ for FQ184 and 8.1 x 10^3^ for FQ151. The data were analyzed by ANOVA to test for differences between strains, lungs and days, and interactions between day*lung and strain*lung. Total bacterial load was significantly different once again between strains (F_2,40_ = 12.0, *p* < 0.001) and was dependent on lung (F_2,40_ = 6.03, *p* = 0.005). There was a main effect of day (F_1,40_ = 155, *p* < 0.001) and the magnitude of the drop between 48 h and 7 days was unaffected by lung (day*lung interaction F_2,40_ = 9.5, *p* = 0.110) and did not vary between strains (strain*day interaction F_2,40_ = 1.73, *p* = 0.190). Lung identity did, however, affect the growth of the strains differently (lung*strain interaction F_4,40_ = 2.62, *p* = 0.049).

**Figure 2.**
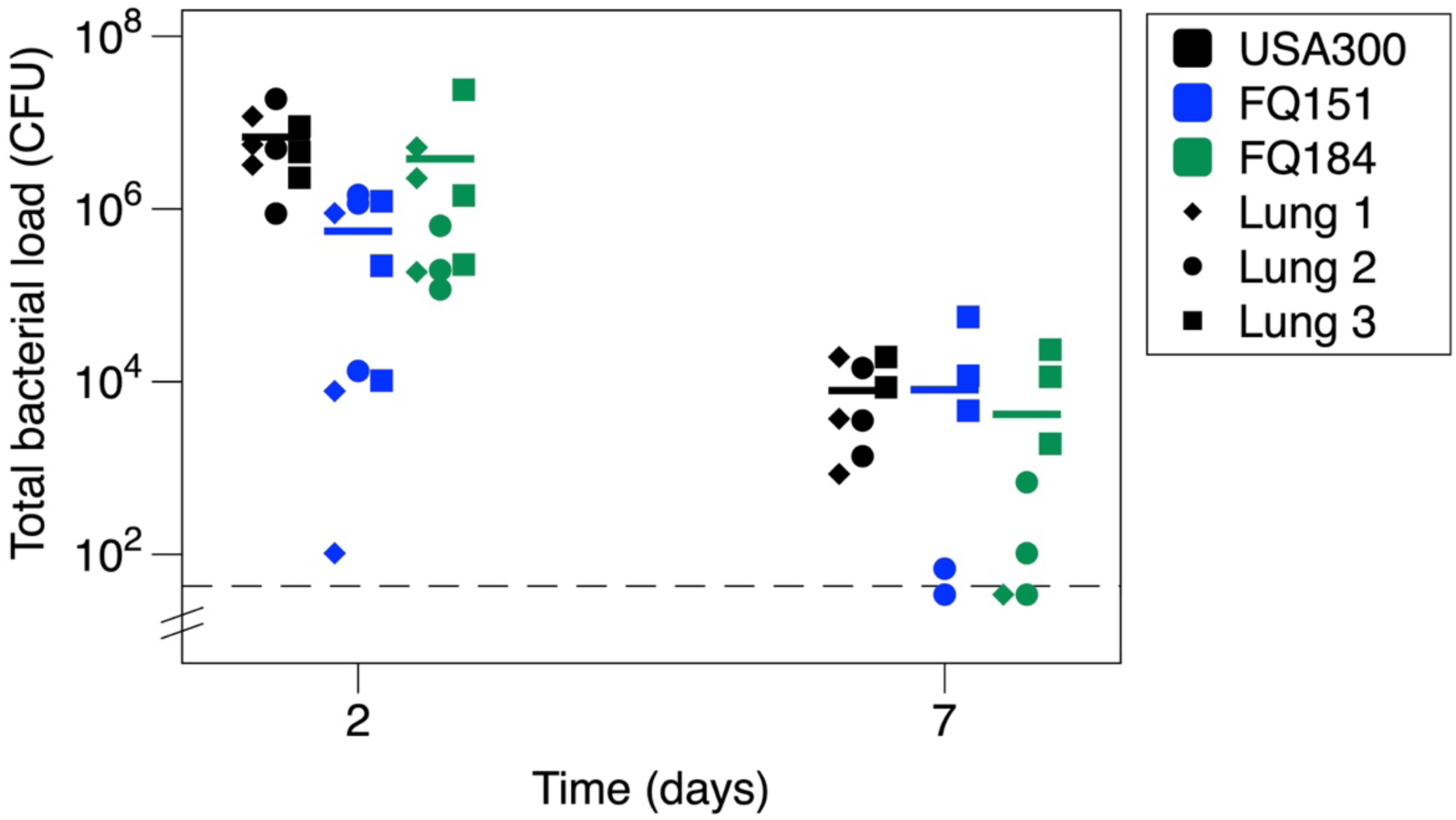
Growth of an exacerbation vs chronic clinical isolate of *Staphylococcus aureus* in the *ex vivo* pig lung model EVPL. *S. aureus* isolates were inoculated into six replica bronchiolar sections of tissue from each of three independent pairs of lungs, and samples were destructively sampled at 48 h and 7 days post infection. Bacterial load was measured as total colony forming units per tissue sample (CFU). Triplicate lungs are indicated by point shape, and lines represent median values. The limit of detection is shown as a dashed line. USA300 is used as a representative control strain. Clinical isolates supplied by the Instituto de Investigación Sanitaria La Fe and numbers are designated as supplied. ANOVA conducted on log-transformed data revealed significant main effects of strain (F_2,40_ = 12.0, *p* < 0.001), lung (F_2,40_ = 6.03, *p* = 0.005) and day, and a significant interaction between lung and strain (F_4,40_ = 2.62, *p* = 0.049). There was no significant interaction between day and lung (F_2,40_ = 9.5, *p* = 0.110) or day and strain (F_2,40_ = 1.73, *p* = 0.190).

### *S. aureus* biofilm is visible at the airway-tissue interface after 7 days incubation

Due to the decline in CFU recovered by day 7 it was important to investigate whether there was any visible *S. aureus* biofilm on the lung or if the model no longer supported growth. Lung tissue was washed in PBS and prepared for microscopy at 7 days post infection. Histopathological staining (Figure 3) showed a clear *S. aureus* biofilm architecture in infected lungs at day 7 which is absent from the uninfected lung. The morphological characteristics of the biofilm appear to vary between strains, but lung tissue remained largely intact across all tested isolates. Although there was some disruption of tissue integrity at the surface (arrows, Fig 3b-c), this was less prominent with the clinical isolates than with USA300. This may reflect these isolates being better adapted to persistence in the lung environment, particularly FQ184 (Fig 3d) (although, interestingly, this was isolated during an acute exacerbation). None of the samples show any evidence of lung abscess, which would present as a clearly-defined, bordered structure present below the tissue surface. The *S. aureus* biofilm is associated with the tissue airway interface and does not appear to demonstrate invasion of the tissue.

**Figure 3.**
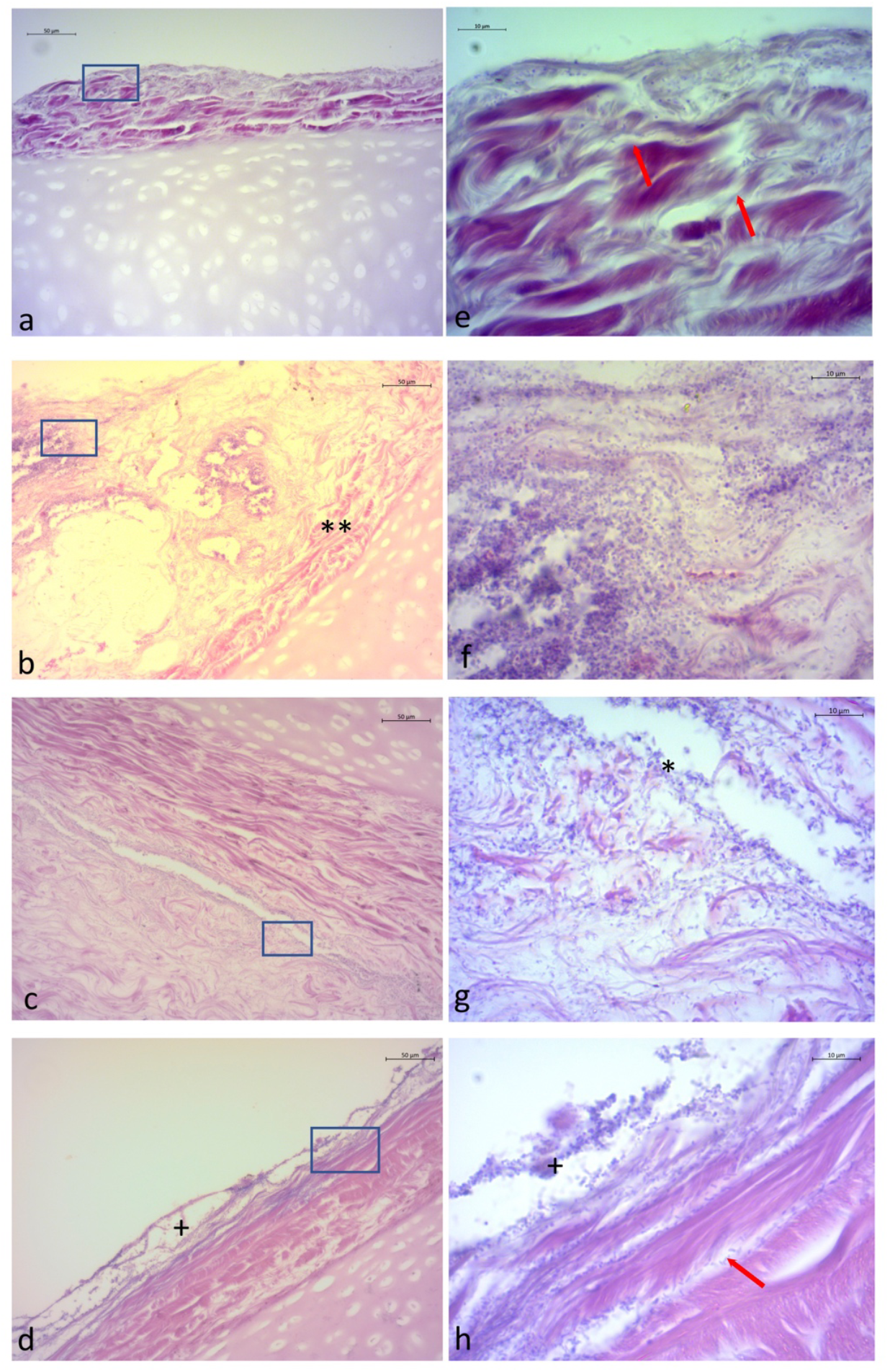
H & E stain of representative lungs at day 7. Uninfected control (a,e) and 7 days post inoculation with USA300 (b,f), FQ151 (c, g) and FQ181 (d, h). 20x magnification (a-d) and highlighted area of interest 100x magnification (e-h). Biofilm architecture is evident at the tissue surface (b and d) and airspace between lung sections (c) but is absent on the uninfected control (a). Clusters of cocci shaped cells are present in all inoculated tissue at 100x (f-h) but not uninfected control (e). Arrows indicate presence of rods and isolated cocci closely associated with tissue (e). Differences in the growth of USA300 (b) and clinical strains (c, d) can be seen. Clinical strains demonstrate less tissue damage (**) closer adherence to the tissue (*), apparently fewer cell numbers but potentially tighter biofilm architecture (+).

There is evidence of other bacterial species in some samples, for example rods can be clearly seen in the tissue infected with FQ184 (red arrow, Fig 3c, 100x magnification). These are likely to be endogenous populations present prior to dissection or contaminants not removed during sterilisation. Interestingly, when other bacteria are present, there appears to be distinct ecological niches within the lung environment with *S. aureus* biofilm growing primarily at or near the surface, and other rods present embedded deeper within the tissue. This supports much of the current understanding of the structural architecture of polymicrobial biofilms and it would be useful to study this further.

### *S. aureus* aggregates in artificial sputum surrounding lung tissue

Initial experiments (Figs 1, 2) only recorded biofilm burden recovered from washed tissue pieces. However, there is evidence in the literature that *S. aureus* may preferentially bind to mucus plugs in the airways of the CF lung, rather than associating with the tissue surface. This could explain why, despite presumably being adapted to the lung environment, CFU loads associated specifically with the bronchiolar tissue showed such a marked drop by day 7. When bacteria were also recovered from the surrounding ASM and enumerated, the total CFU count from the sample (tissue plus surrounding ASM) was higher than the CFU count for tissue only, for all strains except FQ210 (Fig 4a). In fact, by day 7, a greater proportion of the total bacterial population is consistently found in the surrounding ASM than in tissue-associated biofilm (Fig 4b). The four biofilm forming strains all showed a significant increase in the proportion of recoverable cells in the ASM between 48 h and day 7 as measured by the Kruskal Wallis Test. Results were as follows: FQ128 (Chi-squared = 7.44, p =0.006, df = 1), FQ140 (Chi-squared = 11, p =0.0009, df = 1), FQ151 (Chi-squared = 7.1, p =0.008, df = 1) and USA300 (Chi-squared = 12.3, p =0.0005, df = 1). Newman, FQ142 and FQ210 did not show a significant increase in ASM proportion between the two time points (Kruskal Wallis test, Chi-squared = 3, 1.2 and 1.1 and *p* = 0.08, and 0.29 respectively, df =1). As these strains are not biofilm formers (Table 1), this is likely due to a higher proportion already in the ASM by 48 h: for Newman it is clear that growth is not supported in the tissue even by 48 h.

**Figure 4.**
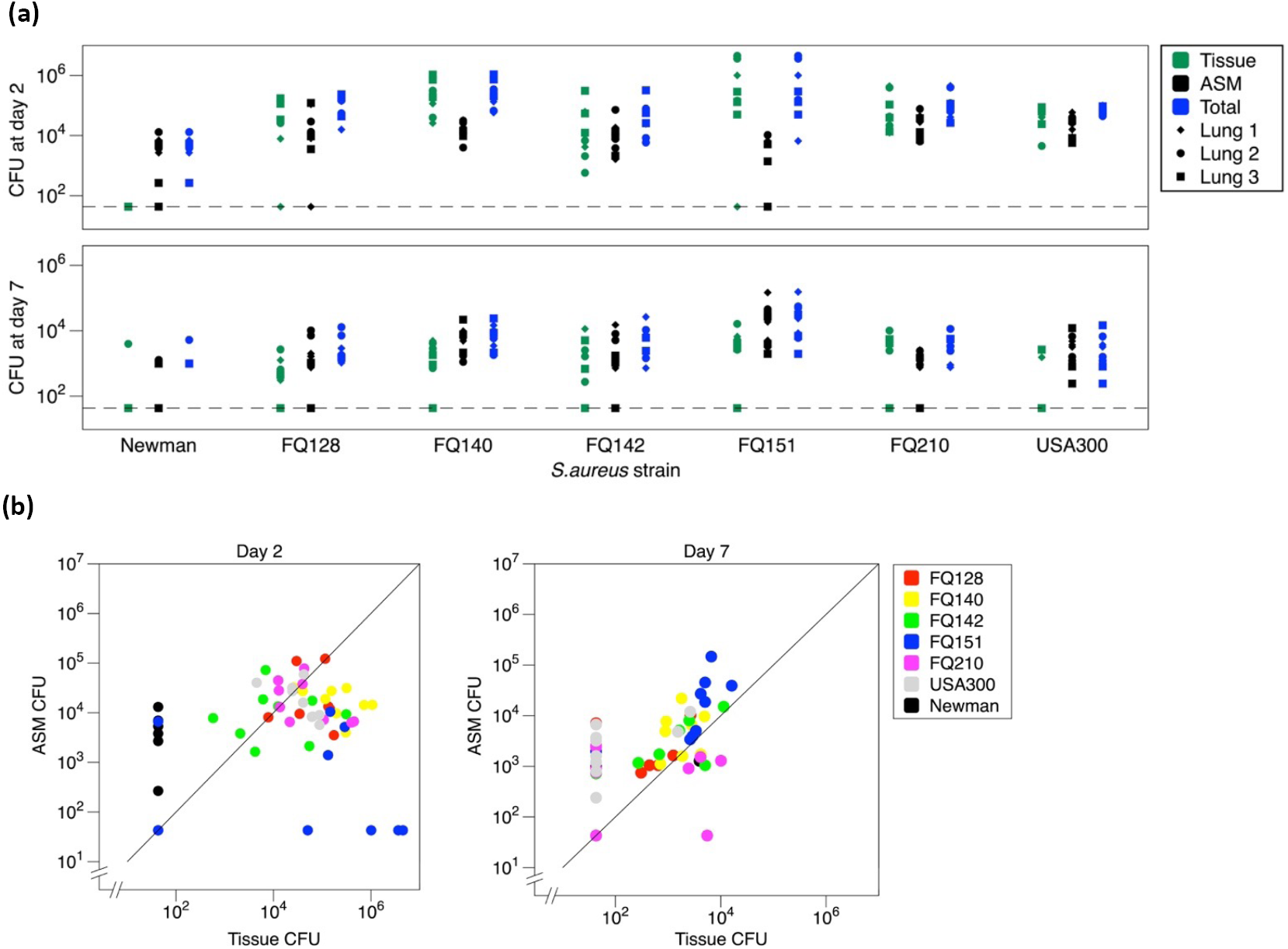
Location of *Staphylococcus aureus* in *ex vivo* pig lung (EVPL) model 48 h and 7 days: CFU and proportion of CFU recovered from Artificial Sputum Media (ASM) vs. tissue. *S. aureus* isolates were inoculated into six replica bronchiolar sections of tissue from each of three independent pairs of lungs, and samples were destructively sampled at 48 h and 7 days post infection. Aliquots of the ASM surrounding each sampled piece of tissue were also assessed for bacterial load. USA300 and Newman were used as representative control strains and an uninfected lung used as a negative control. Clinical isolates supplied by the Instituto de Investigación Sanitaria La Fe and numbers are designated as supplied. **(a)** *S. aureus* cell counts recovered from EPVL at 48 h and 7 days. Bacterial load measured as total colony forming units per tissue, surrounding ASM or total sample (CFU). Counts were taken from triplicate lungs, indicated by point shape. Non-parametric Kruskal Wallis tests were used to test for differences in the proportion of bacteria in the surrounding ASM at 48 h and 7 days. Increases in proportional growth in ASM were found for strains FQ128 (Chi-squared = 7.44, p =0.006, df = 1), FQ140 (Chi-squared = 11, p < 0.001, df = 1), FQ151 (Chi-squared = 7.1, p =0.008, df = 1) and USA300 (Chi-squared = 12.3, p < 0.001, df = 1). Newman, FQ142 and FQ210 showed no significant increase in ASM proportion (Chi-squared = 3, 1.2 and 1.1 and *p* = 0.08, 0.22 and 0.29 respectively, df =1). **(b)** Proportion of total CFU recovered from tissue vs surrounding ASM at 48 h and 7 days. Line shows equal CFU in ASM and tissue, for reference. Minimum limit of detection 4.3 x 10^1^. Linear models on log-log transformed data showed a significant correlation between growth associated with tissue and growth in surrounding ASM at both time points. (Full details of ANOVA results in Supplementary information, R^2^adj for models testing for effects of lung, strain, CFU lung and strain*CFU lung on CFU in ASM were 0.49 for 48 h data and 0.41 for day 7 data; dropping non-significant strain*CFU lung interaction terms showed that the slope of CFU in ASM on CFU associated with lung tissue was <1 at 48 h with *t* = 2.92, *p* = 0.005 but not significantly different from 1 at day 7 with *t =* 1.79, *p* = 0.081).

### Phenotypes consistent with chronic disease are observed within the model

#### a) Small Colony Variants (SCVs)

Small colonies (<50% diameter of usual *S. aureus* colonies) were observed on MSA plates after incubation for at least 48 h in 5% CO_2_ at 37 °C for samples taken 7 days post inoculation. Exemplar colonies were photographed (Fig S1). These potential SCVs were tested for identification and confirmation against WT colonies (Table 2). Despite having weak catalase and coagulase results in some tests, which may be attributed to their small size, SCVs were confirmed as *Staphylococcus* spp. using specific primers (STaG) (38) and as *S. aureus* via sequencing of the 16s-23S intergenic region.

**Fig S1.**
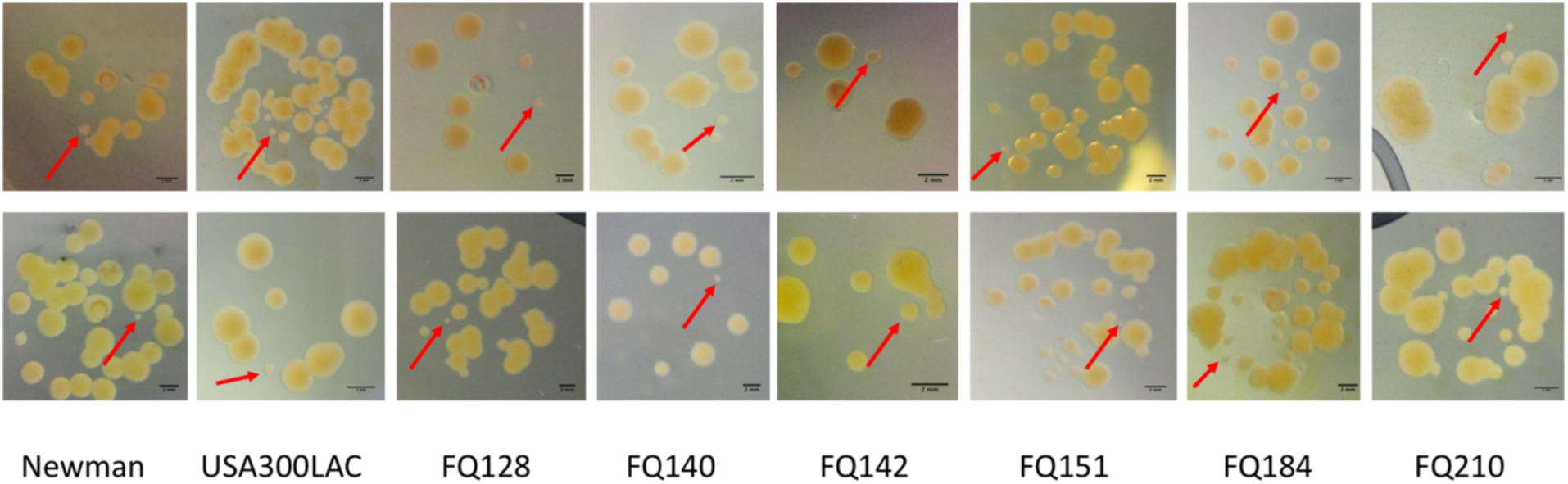
Appearance of Small Colony Varients (SCVs) on MSA plates. Samples taken from lung homogenate (top line) or surrounding Artificial Sputum Media (ASM) (boGom line) at 7 days post inoculation and incubated for 24 h at 37 °C in 5% CO2 and a further 48 h in ambient conditions. Red arrows indicate typical SCV, selected and identified by catalase and coagulase tests and by PCR (Table 1).

**Table 2.** Identification of Small Colony Variants (SCVs). Catalase and coagulase result and colony identity as confirmed by 16S sequence. Samples were taken from lung homogenate or surrounding Artificial Sputum Media (ASM) at 7 days post inoculation and grown on MSA plates for 24 h at 37 °C in 5% CO_2_ and a further 48 h in ambient conditions. SCVs were selected as shown in Fig S1 and all confirmed with *Staphylococcus* specific primers prior to sequencing of the 16S-23S intergenic spacer.

| Strain originally inoculated | Diameter of small colony identified (% WT) | Coagulase | Catalase | 16S sequence identity |
| --- | --- | --- | --- | --- |
| Newman | 42 | + | + | *S. aureus* |
| Newman | 25 | + | + | *S. aureus* |
| USA300 | 27 | + | + | *S. aureus* |
| USA300 | 32 | + | + | *S. aureus* |
| FQ128 | 31 | + | + | *S. aureus* |
| FQ128 | 23 | + | + | *S. aureus* |
| FQ140 | 26 | Weak | + | *S. aureus* |
| FQ140 | 25 | Weak | + | *S. aureus* |
| FQ142 | 43 | + | + | *S. aureus* |
| FQ142 | 43 | + | + | *S. aureus* |
| FQ151 | 40 | Weak | + | *S. aureus* |
| FQ151 | 31 | Weak | + | *S. aureus* |
| FQ184 | 28 | + | + | *S. aureus* |
| FQ184 | 27 | Weak | Weak | *S. aureus* |
| FQ210 | 32 | Weak | + | *S. aureus* |
| FQ210 | 23 | Weak | + | *S. aureus* |

#### b) Antibiotic tolerance

Tolerance to rifampicin was observed for four selected clinical isolates and USA300 when infected tissues were incubated with rifampicin at >500-fold MIC (2 μg mL^-1^) for 24 h at 37 °C (Fig 5). Uninfected control tissues showed no growth on MSA agar plates. All selected *S. aureus* strains showed a sensitive resistance profile to rifampicin, with an MIC of 0.0039 μg ml^-1^ (Table S1). The data were analyzed by ANOVA to test for differences between strains, lungs and rifampicin treatment. The analysis showed an overall decrease in bacterial load across strains following rifampicin exposure (F_1,49_ = 468, *p* < 0.001), and this was not strain dependent (strain*treatment interaction F_1,49_ = 1.81, *p* = 0.141). Consistent with biofilm growth conferring antibiotic tolerance, only 2-3 log killing was found on exposure to >500-fold MIC.

**Figure 5.**
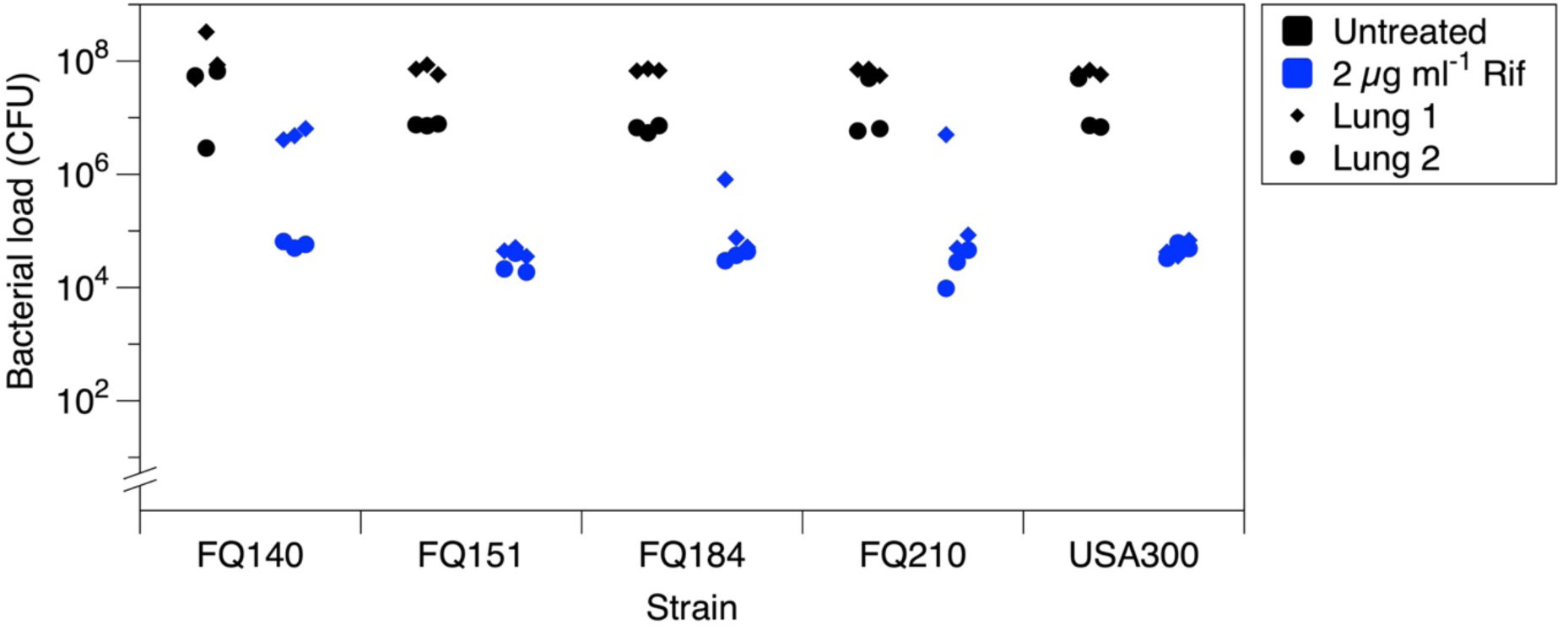
Antibiotic tolerance in the *ex vivo* pig lung model (EVPL) model. Each *S.s aureus* isolate was inoculated into six replica bronchiolar sections of tissue from each of two independent pairs of lungs, and incubated as previously described for 48 h. Three infected replica were washed, homogenized and plated for counting of bacterial load, while the other three infected replica were moved to MHB containing rifampicin at >500-fold MIC (2 μg ml^-1^) for 24 h before being washed, homogenized and plated. Six uninfected tissue pieces were used as control and processed as infected replica. All isolates were classified as sensitive by standard antibiotic susceptibility testing. The results demonstrated a non-significant decrease in bacterial load of 2-3 log, which demonstrates antibiotic tolerance in the EVPL model. Data were anlayzed by ANOVA and this showed an effect of lung on overall bacterial load (F_1,49_ = 46.6, *p* < 0.001), differences between strains in total bacterial load (F_4,49_ = 5.62, *p* = 0.001) and a consistent decrease in bacterial count following rifampicin treatment (treatment F_1,49_ = 468, *p* < 0.001; strain*treatment interaction F_1,49_ = 1.81, *p* = 0.141).

## Discussion

We have shown that *S. aureus* infection can be established in an *ex vivo* pig lung model of CF. Results using a biobank of clinical isolates demonstrate that, in a single lung, strains grew with reasonable consistency as tissue-associated biofilm and as bacterial aggregates in artificial mucus. Bacterial bioburden was maintained over 7 days, although biofilm numbers were diminished over this period. The EVPL model is high throughput and inexpensive; also, because it uses post-consumer waste from the meat industry, it avoids the ethical concerns and limitations inherent in live animal models (35). Several results from this work demonstrate that the model (a) cues *S. aureus* to express phenotypes characteristic of chronic infection, and (b) recapitulates key aspects of the aetiology of *S. aureus* in chronic CF which are not captured by the current “gold standard” mouse models of respiratory infection. We will now discuss the strengths and limitations of our model.

### Chronic infection phenotypes of *S. aureus* in EVPL

In EVPL, *S. aureus* isolates exhibited enhanced tolerance to rifampicin, likely due to a combination of physiological cues from the lung environment, the formation of biofilm and the appearance of a subpopulation of SCVs (10, 12, 40-42). The phenotypic development of SCVs has been linked to chronic CF infections as it permits longer survival of the bacterial cells (40, 41). Extensive biofilms typical of *P. aeruginosa* have not been described for *S. aureus* in the CF lung (43), but biofilm-like aggregates are reported in the sputum and on the surface of airways (23, 44-46). Consistent with growth as biofilm aggregates, there is evidence that *S. aureus* switches off the global regulator *agr* in CF (47). Generally, interstitial bacteria are rarely observed in biopsy samples from people with CF (even if significant interstitial inflammation is observed) and bacteria in the airways are confined in luminal mucus plugs or occasionally attached to defined foci of epithelial erosion; this lack of tissue invasion is consistent with the rarity of bacteraemia in people with CF (23, 45, 48-53). CF lung disease thus presents as bronchiectasis with mucous plugging of small airways. In contrast, microscopy images presented in studies of murine pulmonary *S. aureus* infections typically show abscesses (a cavitating, pus-filled lesion within the tissue with a defined border) (see Figure 2 in ref. (28) and Figure 2 in ref. (54) for example) and lead to chronic pneumonia (55). Clearly, there are significant differences in *S. aureus* pathology in mouse models *versus* the highly specialized environment of CF airways.

In light of this, it is significant that our results suggest preferential localisation of *S. aureus* in the ASM surrounding the tissue sections, as opposed to epithelial surface attachment, and do not lead to the appearance of abscess-like structures. *S. aureus* can cause lung abscess in humans – in fact it is one of the most common bacteria isolated from abscesses (56) – but abscesses are almost never observed in people with CF (52, 53). A particular Panton-Valentine Leukocidin (PVL)+ strain of MRSA was associated with invasive, cavitating lung lesions in a CF centre in the USA in the mid 2000s (57). Other than this a literature search revealed only five cases of abscess among people with CF, only two of which were associated with *S. aureus*: one patient was co-colonized by *P. aeruginosa* and the other with *P. denitrificans* (31, 58). The rarity of clinical evidence for invasive *S. aureus* infection in people with CF also calls into question the likelihood of intracellular invasion by this species, even though it has been documented in epithelial cell cultures *in vitro* (59, 60). The clinical observations are, however, entirely consistent with a study in which mucus hypersecretion was induced in cultured primary nasal epithelia cells to better mimic conditions in the CF lung: mucus presence led to *S. aureus* cells moving away from the epithelial cell surface and growing in the mucus (23) – as observed in our model.

### Population heterogeneity of *S. aureus in vivo* and in EVPL

The evidence outlined above is consistent with *S. aureus* potentially growing as phenotypically heterogeneous populations both in CF and in EVPL – comprising tissue-associated biofilm aggregates and mucus-embedded aggregates, and a proportion of SCVs. Persistence, antibiotic tolerance, gene expression and ability to culture *in vitro* are likely to differ between these subpopulations, and understanding the differences within heterogenous populations could improve understanding of virulence mechanisms. If this is the case, the model will be valuable in future study for the generation of heterogenous populations that are hard to reproduce in animal and traditional *in vitro* models.

### Strengths and limitations of EVPL as a model for *S. aureus* CF infection

The use of *in vivo* animal models is limited by ethical concerns, cost and host immune response (35). In an effort to reduce and replace the use of animal models, *in vitro* models have been developed to represent the molecular mechanisms and phenotypic characteristics of disease. Tissue culture monolayers, using cell lines and primary cell culture, have been used extensively to study the interaction of the host epithelial layer and pathogens, most notably *P. aeruginosa* (61-66). However, these models are often over simplistic in nature and lack properties relating to the host tissue such as mucus hypersecretion, cell polarity, differential cell types, and tissue structure and integrity (67). Biofilms in these models were grown on abiotic surfaces and such *in vitro* biofilms have been shown to differ from *in vivo* biofilm in a number of significant ways (68). More specifically for the incidence of *S. aureus* in CF, it is also abundantly clear that both mouse and *in vitro* models fail to capture key features of pathology observed from human clinical data. Our EVPL model could thus fill an important gap in our toolkit for answering crucial questions about the microbiology and clinical impact of *S. aureus* in CF. To this end, we present a critique of the model as it currently stands.

Variability in *S. aureus* bacterial load in the model is considerable, and there are a number of possible explanations. For example, there may be differences in starting inocula. Lungs were inoculated from single colonies, and not a standardized broth culture, as preliminary work with *P. aeruginosa* in the model showed no significant variation in the number of cells inoculated from single colonies, or the cell numbers recorded at 48 h (34). However, given the differences in the way that the two bacteria grow in the model, starting inoculum may be a more significant determinant of the growth and survival of *S. aureus* in the model thus warranting further study. Standardized cell numbers would be important if, for example, the model were to be developed to measure the impact of antimicrobial agents on viable bacterial burden. It is worth noting that if the tissue-associated subpopulation of *S. aureus* grows as a biofilm and not within the tissue (Fig 3), then surface area of tissue sections may have a significant impact on the bioburden in each tissue sample and further standardisation of tissue sample size and shape may improve consistency.

Conversely, in the lung airway there is unlikely to be a large burden of colonising bacteria (69), but rather a few seeding cells transferred from the upper airway. In this respect, inoculating from a single colony may actually more closely reflect the clinical situation, and was therefore regarded as sufficient to study the initial establishment, growth and maintenance of *S. aureus* within the scope of this study. More data on the variability in *S. aureus* aggregate size between foci of infection within the lungs, and between patients, would be needed to answer this question. In a study by Hirschhausen *et al.* (55), adaptive changes differed in patients infected with the same *S. aureus* clonal lineage, indicating that individual host factors had an impact on adaption. This is reminiscent of our results, which show significant variability between lungs taken from different pigs (Fig 2).

In our initial growth studies (Figures 1 and 2) the decline in CFU count between day 2 and 7 was consistent, suggesting that despite variation between strains and lungs, growth patterns for the same strain were reproducible. The reduction in cell count was also not due to the model being unable to support *S. aureus* growth, as *S. aureus* CFU were observed at 7 days and this was confirmed histologically (Fig 3). Differences in the growth and localisation of a pair of isolates taken from the same patient during stable infection and acute exacerbation also warrant further exploration with a larger collection of isolates (Fig 3).

Further empirical work, conducted with close attention to the available clinical data, is required to fully assess the strengths and limitations of EVPL as a model for *S. aureus* CF infection. However, the data we present show striking consistencies with clinical observations which are not matched by mouse models, and back up the results of the only cell line study which mimicked CF conditions by inducing mucus hypersecretion. We thus conclude that EVPL plus ASM could be a valuable addition to the suite of infection and biofilm models available to microbiologists.

## Conclusion

Our results confirm that *S. aureus* infection of an *ex vivo* model comprising pig bronchiole and artificial cystic fibrosis mucus demonstrates key similarities with chronic CF infection: aggregation of bacterial cells in tissue-associated biofilm but also in mucus; the development of SCVs; and an increase in antibiotic tolerance. The model is also able to distinguish differences in growth between clinical strains. The potential preferential binding of *S. aureus* to mucus, and lack of tissue invasion, is a phenomenon that may have clinical relevance that is not observed in current murine models of pulmonary infection.

## Acknowledgements

We would like to acknowledge the help of Cerith Harries and Caroline Stewart and the use of the media preparation facilities within the School of Life Sciences technical team, and Ian Hands-Portman in the Life Sciences imaging suite, at the University of Warwick. We also thank Peter Walker and the Histology Core Facility at the University of Manchester for histological sample preparation. Finally, we thank Amparo Soler at Hospital Universitari i Politecnic La Fe, for the demographics and clinical presentation data of CF patients. The work was funded by a Medical Research Council New Investigator Research Grant to FH.

## Author Contributions

FH & ES conceived the study. ES conducted experimental work, analyzed data and drafted the manuscript. FH developed the EVPL model. MMH conducted experimental work and analyzed data (rifampicin tolerance) and contributed to manuscript preparation. NEH conducted experimental work (histological sample preparation and imaging), aided with data analysis and contributed to manuscript preparation. MATM contributed clinical isolates and associated data. ARS and MNH contributed clinical context for the study goals and interpretations, and contributed to manuscript preparation. All authors saw and approved the final manuscript draft.

